# GradHC: Highly Reliable Gradual Hash-based Clustering for DNA Storage Systems

**DOI:** 10.1101/2023.10.05.561008

**Authors:** Dvir Ben Shabat, Adar Hadad, Avital Boruchovsky, Eitan Yaakobi

**Affiliations:** Computer Science Department, Technion, Haifa, 3200003, Israel

## Abstract

As data storage challenges grow and existing technologies approach their limits, synthetic DNA emerges as a promising storage solution due to its remarkable density and durability advantages. While cost remains a concern, emerging sequencing and synthetic technologies aim to mitigate it, yet introduce challenges such as errors in the storage and retrieval process. One crucial in a DNA storage system is clustering numerous DNA reads into groups that represent the original input strands. In this paper, we review different methods for evaluating clustering algorithms and introduce a novel clustering algorithm for DNA storage systems, named Gradual Hash-based clustering (GradHC). The primary strength of GradHC lies in its capability to cluster with excellent accuracy various types of designs, including varying strand lengths, cluster sizes (including extremely small clusters), and different error ranges. Benchmark analysis demonstrates that GradHC is significantly more stable and robust than other clustering algorithms previously proposed for DNA storage, while also producing highly reliable clustering results.

## 1 Motivation

DNA storage is an emerging technology that promises to revolutionize the way we store and preserve digital data [1–3]. By encoding data as synthetic DNA, it becomes possible to store vast amounts of information in a very compact form that can last for thousands of years, without the need for electricity or any other external power source [3,4]. Moreover, DNA is much more durable than other storage media, such as hard drives, which can degrade or fail over time, and it is also very energy efficient [1, 3, 5]. However, the cost of DNA storage is still a significant barrier to its widespread adoption. Although the price of DNA synthesis has been declining rapidly in recent years, it is still much higher than other storage technologies [1–5]. Moreover, the process of storing and retrieving DNA data is complex and time-consuming, involving several steps that require specialized expertise and equipment.

To use DNA as a storage medium, the first step is to map binary data onto DNA strands by replacing every two bits with their corresponding DNA nucleotide (adenine (A), thymine (T), guanine (G), and cytosine (C)). Because of limitations in synthesis technology, the data is divided into short strands no longer than a few hundred nucleotides [6,7]. Next, these DNA strands can be encoded using error correcting codes or other encoding techniques. These DNA strands are then synthesized, yielding hundreds to thousands of noisy copies of each original strands which are stored in a container. When data needs to be retrieved, a small sample of DNA strands is taken from the container and amplified using a technique called polymerase chain re-action (PCR). After amplification using PCR, each DNA strand is sequenced to determine its DNA symbols. However, the synthesis, PCR, and sequencing steps are all susceptible to errors, including insertions, deletions, and substitutions, which can result in errors in the stored data. Each error type occurs with a different probability, and the overall error rates are mainly influenced by the synthesis and sequencing technologies used [2–4, 8]. The outcome of the sequencing step is an unordered set of thousands to millions of DNA strands, consisting of multiple noisy copies of the original DNA strand. Before the decoding process, a *clustering algorithm*, which is the focus of this work, must be used to group the noisy copies together so that each cluster contains copies of the same original DNA strand. After clustering, each cluster is reconstructed using a *reconstruction algorithm* to reconstruct an approximation of the original DNA strand [9–13]. Finally, once all the DNA strands have been retrieved, the coding scheme is used to decode them, and the reverse mapping is applied to obtain the original binary data.

In this work, we aim to address the problem of clustering in DNA-based storage systems. We will review all existing clustering algorithms and identify their strengths and weaknesses. In addition, we will propose the GradHC algorithm, a novel clustering approach specifically designed for DNA storage systems. To evaluate the effectiveness of our proposed method, we will conduct benchmarking experiments on previous experiments and simulated datasets and compare the results against those of state-of-the-art clustering algorithms for DNA storage.

## 2 DNA Clustering

### 2.1 Problem Definition

As presented in the Introduction, a key aspect of every DNA data-storage-system is the *clustering* phase. When reading the strands after the sequencing process, one gets multiple noisy copies from every input strand. The number of copies vary from a few to thousand’s and depends on the sequencing technology, the number of sampling cycles, and the PCR process. Before applying a *reconstruction* algorithm to get the original input, one must cluster the strands together, in a way that with high probability every cluster consists of noisy copies of the same input strand. Formally, a clustering *C* of a finite set *S* ⊆ Σ^∗^, is any partition of *S* into nonempty subsets, when Σ = {*A, C, G, T}* and *S* = {*s*_1_, *s*_2_, …, *s*_*n*_} is the set of all DNA reads gathered in the sequencing process. One must note that clustering DNA strands differ from traditional clustering problems in the way that the clustering algorithm is not given with essential parameters settings such as density threshold or cluster size distribution. In addition, computing a distance matrix between every pair of input strands is necessary for some clustering algorithms, but it is not feasible due to the large scale input size of DNA-based storage systems. The edit distance^1^ algorithm, which is commonly used for computing string distances, has a quadratic time complexity of *O*(*l*^2^), where *l* is the length of the input strands. This means that as the input size increases, the computational cost of computing the edit distance matrix grows exponentially. Thus, making the clustering task of DNA-storage system data is a much more difficult problem. Moreover, in DNA storage systems, strands are not ordered in the memory, making it challenging to determine their storage order. The solution is to use indices that are stored as part of the strand to indicate its location relative to other strands. However, a naive algorithm that solely uses the index to cluster the strands may not work well due to potential errors in the index field.

### 2.2 Related Work

There are several DNA-clustering algorithms, where most of them are from the field of bioinformatics and metagenomics. UCLUST [14] and CD-HIT [15] are among the most used clustering methods, both of them use greedy algorithms for the clustering process while they rely on the Needleman-Wunsch global alignment algorithm for calculating the similarity between the sequences. DNACLUST [16] is another greedy clustering algorithm, which uses fast sequence alignment techniques and k-mer based filtering. MeShClust v1.0 [17] uses the famous mean shift algorithm in order to overcome the challenge that when using a greedy algorithm there is no guarantee that an optimal solution can be found. MeShClust v3.0 [18] uses the mean shift algorithm and alignment-free identity scores for the sake of not paying precious time due to the costly global alignment algorithm. MetaDEC [19] applies a deep unsupervised learning approach to cluster metagenomic DNA strands. MM-seqs2 [20] uses a graph-based approach, in which every noisy copy is a vertex, and two noisy copies are connected if they satisfy a particular similarity criteria.

One main weakness of some DNA-clustering algorithms, including those mentioned above, is that they rely on a single sequence identity threshold applied to every cluster. This biologically comprehensible parameter ranges from 0 (no sequence similarity) to 1 (identical strands). However, selecting an inappropriate identity threshold can lead to poor quality clusters. Determining the optimal threshold value requires users to have domain knowledge, which can be complex in DNA storage systems. ALFATClust [21] uses rapid pairwise alignment-free distance calculations and community detection to generate clusters, dynamically determining the cut-off threshold for each individual cluster based on cluster separation and intra-cluster sequence similarity. SEED [22] uses advanced hashing techniques and interval seeding to index DNA strands.

All of the algorithms mentioned above are designed for clustering genomic or meta-genomic data with the goal of grouping similar DNA strands into clusters, which can then be assembled. In DNA storage systems, the clustering step aims to divide the noisy copies so that each cluster can be independently reconstructed afterwards. A key difference is that in DNA storage systems, the DNA strands are much shorter (50-200 nucleotides) compared to genomic data, which consists of long reads of tens of thousands of nucleotides. Therefore, these algorithms are not suitable for clustering short DNA strands.

Starcode [23] clusters the noisy copies based on all-pairs search where it derives the Levenshtein distance^2^ mainly by using the so-called edit matrices. This algorithm is designed to cluster DNA barcodes - very short DNA strands (tens of nucleotides at most). DBSCAN [24] is one of the earliest clustering algorithms, suggesting clustering together data samples based on their density and identifies outliers as noise. However, Guanjin et al. [25] have shown that this algorithm (as well as other density-based clustering algorithms) is impractical for clustering hundreds of thousands of noisy copies due to computational constraints. Shinkar et al. [26] proposed an index-based clustering algorithm and introduced a novel coding scheme called clustering-correcting codes, which uses both the data and index fields for the encoding process. Nonetheless, the success of this algorithm is guaranteed only under the strong assumption that the majority of strands in every generated cluster are derived from the same original strand. Another DNA clustering approach was presented in a recent work done in [27].

The authors presented a novel distributed algorithm that managed to cluster billions of reads under one hour. The algorithm attractively merges clusters based on random representatives while using a hashing scheme to estimate the edit distance. Antkowiak et al. [28] demonstrated a DNA storage system that relies on massively parallel light-directed highly-error prone synthesis. In their work, they presented an LSH-based (locally sensitive hashing) clustering algorithm. This group managed to successfully store 100 KB of information. Clover is one of the newest DNA storage clustering algorithms available, developed by Guanjin et al. [25]. This algorithm has a linear computational complexity and low memory consumption. It achieves its speed by avoiding computation of the Levenshtein distance and using a tree structure for interval-specific retrieval instead.

### 2.3 Performance Validation

Cluster validation is a computational task, used for evaluation of clustering algorithms in many fields, including the bioinformatics, machine learning, data compression and DNA storage. Currently, a wide variety of clustering evaluation metrics exists in these different fields. The reason for of such a variety is due to the multiple aspects of correctness by which the accuracy and goodness of clustering algorithms can be evaluated, and the different requirements from the clustering results in each field. In most common metrics, clustering results are evaluated based on “ground truth” labels - i.e., perfect clustering labels. The perfect clustering of a DNA storage system is given by clustering every noisy copy with its original strand from which it was created. When simulated datasets are used, the perfect clustering is known in advance, since for each noisy copy generated, the original strand to which it belongs is known during the dataset creation process. Computing the perfect clustering of a wet experiment dataset is a more complex task. The best approach for solving this challenge is through an iterative greedy method: every read is associated with an original strand, in which it has the best edit-distance score with, among all original strands. In this section we will present two popular metrics which will be used to compare between the results of the different clustering algorithms in our work.

#### 1. Threat Score (TS)

the threat score (also known as critical success index (CSI) or Jaccard index [22]) is a verification measure used to quantify the similarity between two datasets. Given a clustering 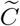 and a perfect clustering *C*, we define *TP, FP* and *FN*, (the overall number of true-positives, false-positives and false-negatives, respectively) by:

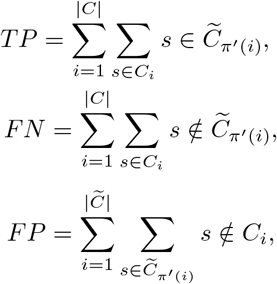

where *π*′ is an injective map

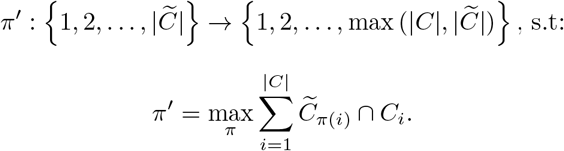

The threat score index takes on a value between 0 and 1, and is defined by the following formula:

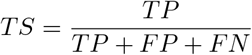

#### 2. Accuracy(*γ*)

This metric was first defined in [27]. Given two clustering 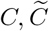, the Accuracy(*γ*) of *C* with respect to 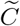, is given by:

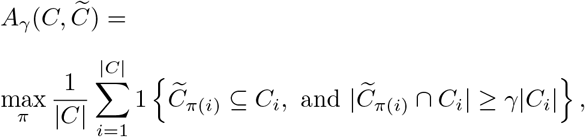

where the maximum is taken over all injective maps:

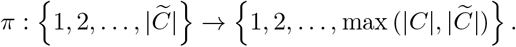

The accuracy 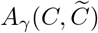 measures the number of clusters in *C* that overlap with some cluster in 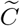 in at least a *γ*-fraction of elements while containing no false positives. It is important to note that the hard constraint that does not allow the clustering algorithm to have false-positives at all, is not necessarily needed in DNA storage systems. This follows since some reconstruction algorithms may overcome the presence of small number of false-positives, if the cluster is sufficiently large. Nevertheless, it should be remembered that in DNA storage systems the sizes of the clusters vary and in many cases may be extremely small. In addition, there is a great significance in evaluation of the clustering algorithm’s results, regardless of the performance quality of the reconstruction algorithm used.

Another clustering evaluation metric was presented in [25]. The authors measures the accuracy of two clustering *C* and 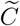 as:

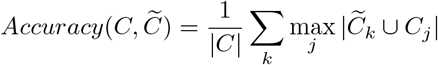

where *C* = (*C*_1_, *C*_2_, …, *C*_*j*_) refers to the perfect clustering and 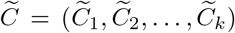 is the output of their clustering algorithm. One can notice that a naive clustering algorithm, which clusters every noisy copy in its own cluster, gets a maximum score for the above metric. For this reason, we chose not to use this metric as part of our performance validation. Other common metrics for clustering algorithm evaluation are *Purity* and *NMI* [21]. *NMI* assumes balanced data point distribution, making it unsuitable for DNA storage systems with unevenly distributed cluster sizes. *Purity* does not penalize misclassifications within clusters and favors clustering solutions with a larger number of clusters, even if they are not meaningful. We chose the metrics listed above, as we find them to be the most suitable and precise for validation of clustering algorithms for DNA storage systems. The *Accuracy(γ)* metric allows to get a decisive indication of the percentage of clusters that managed to restore, so that a good reconstruction algorithm will succeed in reconstructing the original strand from them. This is due to the fact that they contain most of the original cluster, and do not contain noisy copies of other input strand. According to the analysis in [13], reconstruction algorithms yield significantly poor results for clusters that are either too small or too noisy. Therefore, it can be inferred that low *Accuracy(γ)* values of a clustering algorithm directly lead to inaccurate/unfeasible reconstruction of certain clusters in the dataset, negatively impacting the reconstructability of the original data. The *TS* metric allows to get a high level perspective on the full clustering result. In general, a high *TS* value implies two important conclusions: 1. The amount of clusters recovered is high - considering the relatively small number of false-negatives. 2. The generated clusters were recovered with high quality, since the number of false-positives is low compared to the amount of true-positives.

## 3. The GradHC algorithm

### 3.1 Algorithm Description

In this section, we present the *Gradual Hash-based Clustering* algorithm. Our algorithm was developed to deal with a wide range of error rates, and is inspired by the LSH-based clustering, suggested by Antkowiak et al. [28]. We designed an algorithm that can handle datasets with varying clusters sizes, from small (a few strands only) to large (hundreds to thousands of strands). Our clustering algorithm consists of three parts. First, we perform a primary step to obtain a coarse partition of the noisy copies with the aim of minimizing time costs afterward. Next, we execute the second step individually on each chunk of noisy copies generated in the previous step. Finally, a more computationally intensive third step is carried out, taking into account the entire input dataset.

#### 3.1.1. Step 1 - Division into Chunks

##### Algorithm 1

Chunks Generation

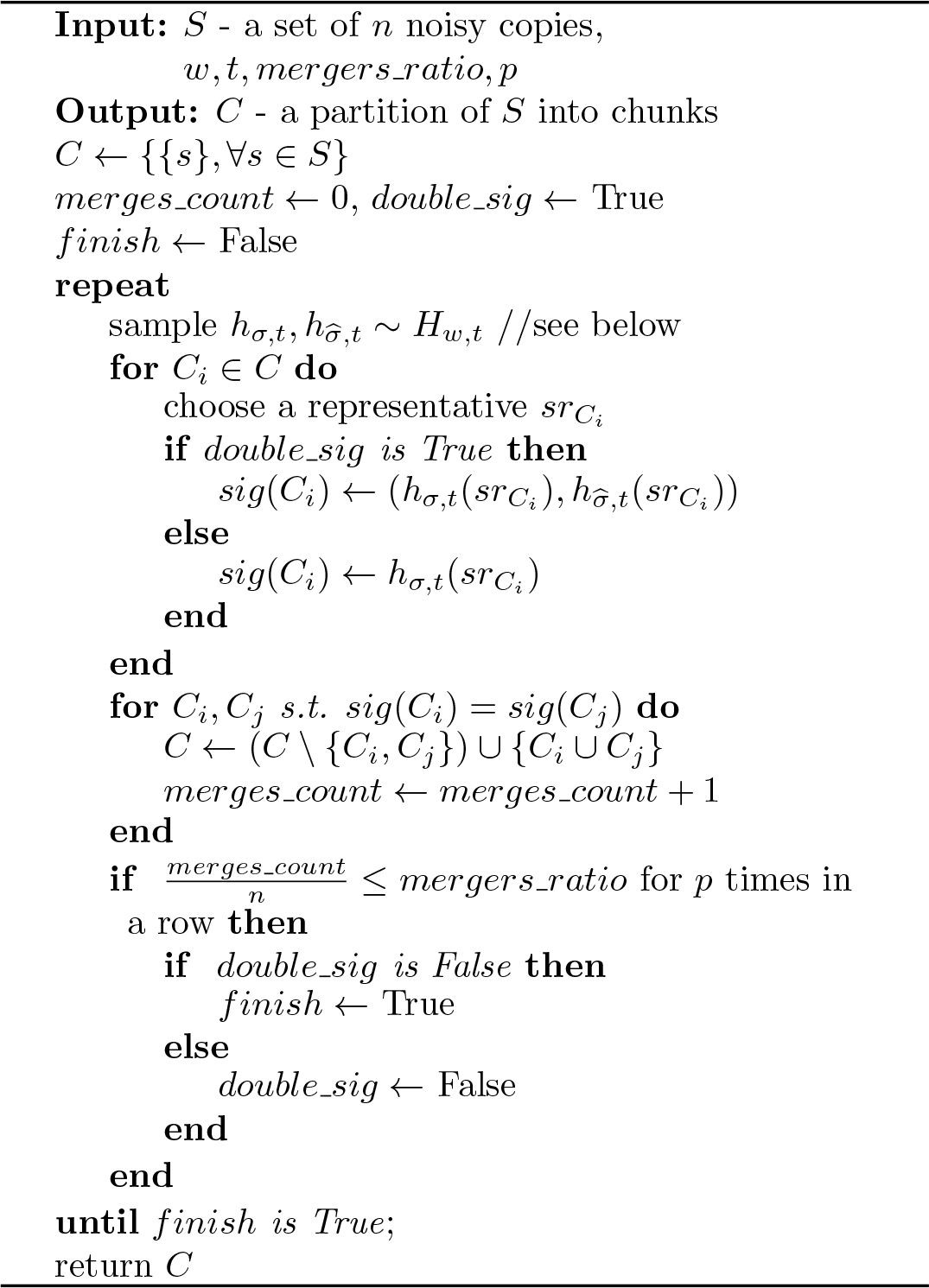

The algorithm begins by dividing the *n* noisy copies of *S* into initial clusters (chunks). The goal is to improve performance in the next stages of the algorithm, where the actual clustering process happens. The partitioning is supposed to be coarse and cheap. Let *x* = *x*_1_*x*_2_…*x*_*r*_ be a given strand, and let *i* be the index of the first occurrence of *y* ∈ Σ^*w*^ in *x* under a given permutation *Σ* : Σ^*w*^ → Σ^*w*^. We define *h*_*Σ,t*_(*x*) to be the substring *x*_*i*_*x*_*i*+1_…*x*_*r′*_, where *r*′ = min(*r, i* + *w* + *t*). As defined in [27], the family of hash functions *H*_*w,t*_, which is parameterized by *w* and *t*, is given by: *H*_*w,t*_ = {*h*_*Σ,t*_ : Σ^∗^→ Σ^*w*+*t*^ }, where *Σ* is a permutation of Σ^*w*^. In the algorithm, we sample *h*_*Σ,t*_ from *H*_*w,t*_ by choosing a uniformly random permutation *Σ*. In this step, we make use of a skeleton of the algorithm suggested by Rashtchian et al. [27], with several modifications. First, as accuracy is not a necessity, we refrain from computing edit-distance. Second, in the first iterations, we use two signatures (generated by their proposed family of hash function *H*_*w,t*_), and only after several iterations the algorithm moves towards relying on a single signature. The motivation behind the changes was to keep the computation fast, resulting with a partition into chunks aimed to resemble the original clusters, making the next steps easier. We set the values for *w* and *t* to be *w* = ⌜log_4_ *l*⌝, and *t* = log_4_ *n* −*w*, as recommended in [27]. The values for *mergers ratio* and *p* (number of ineffective rounds) were chosen empirically and set to 0.0002 and 3, respectively.

#### 3.1.2. Step 2 - Clustering per Chunk

##### Algorithm 2

Clustering Per Chunk

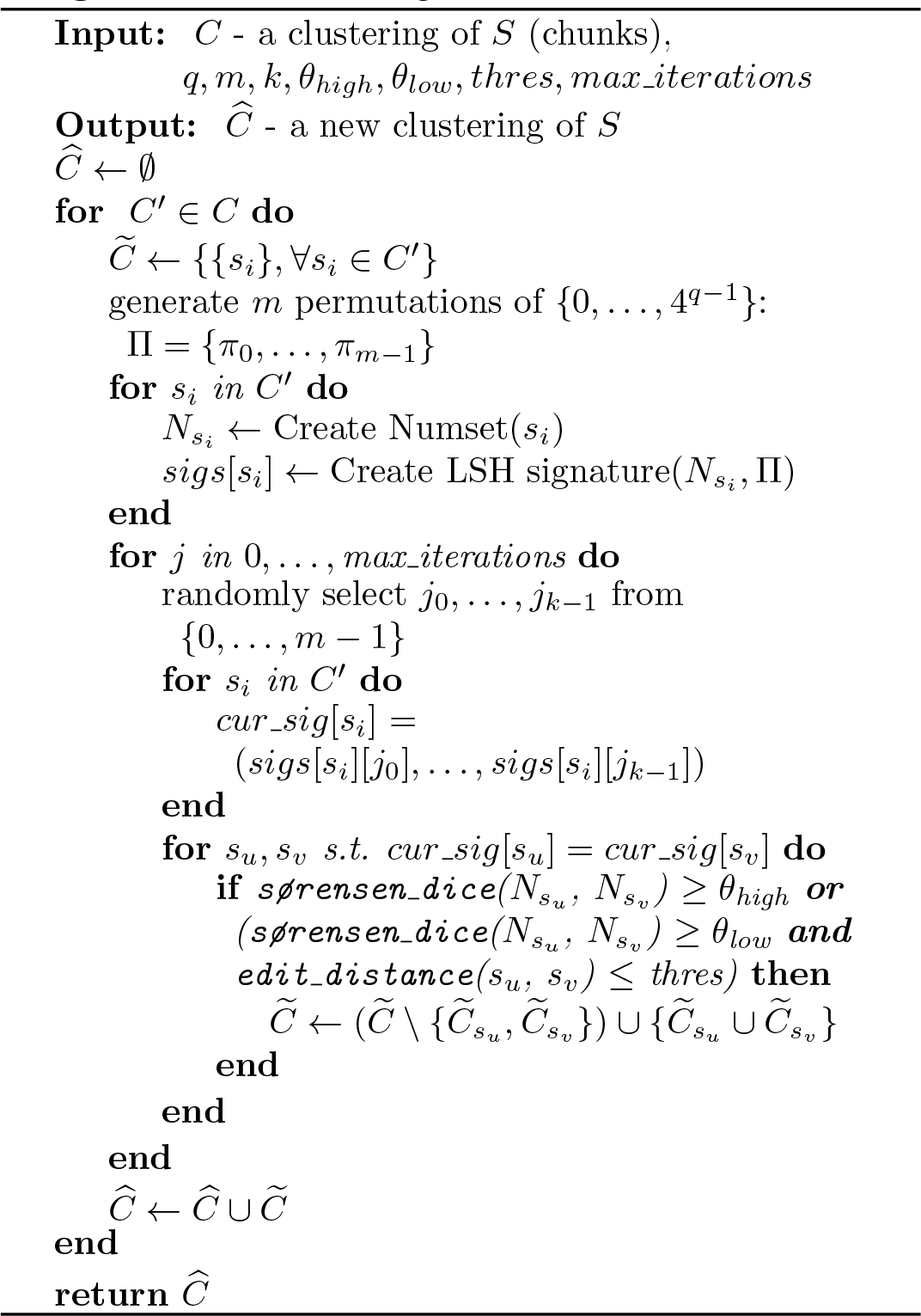

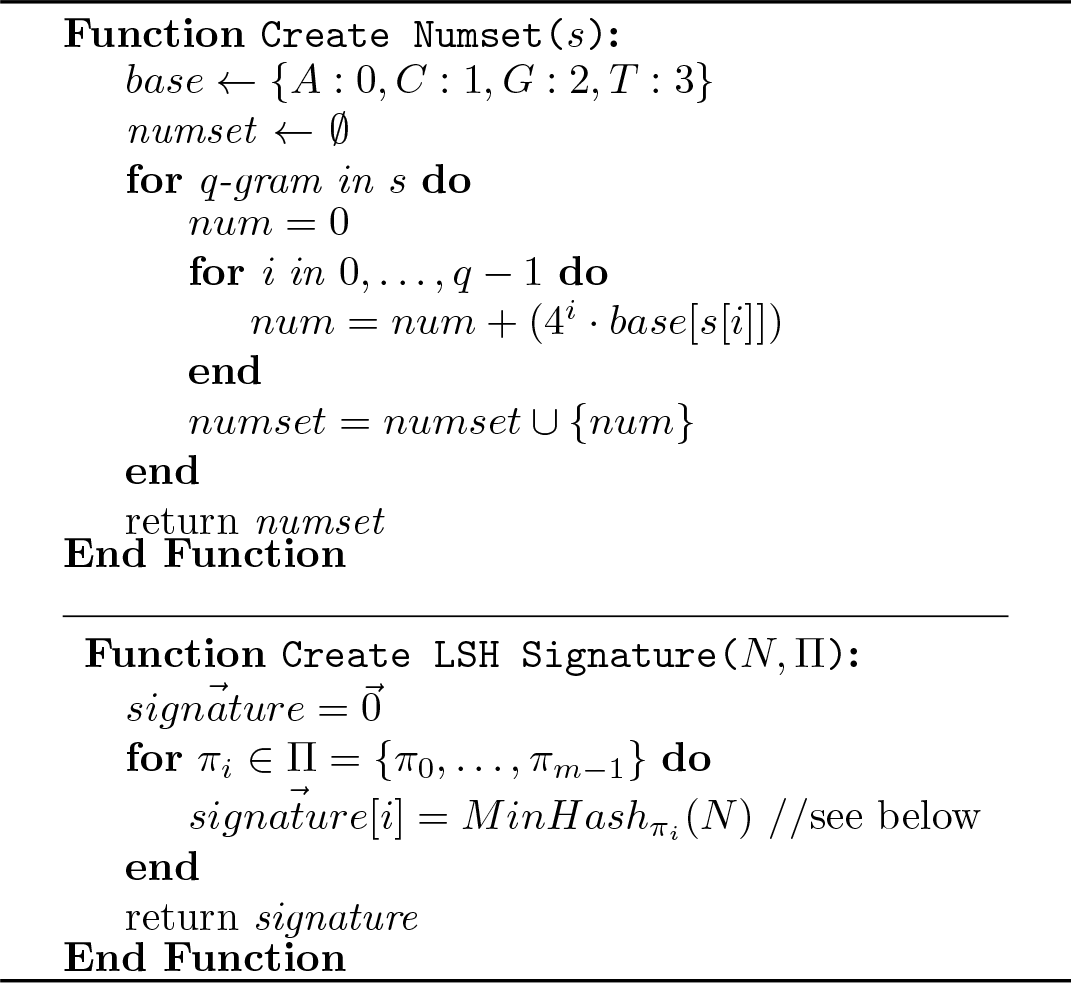

In the second step, the DNA sequences are treated as sets of *q-grams* - overlapping sub-strings of length *q*. By assigning each character from {*A, G, C, T*} a number, we obtain an integer in base 4, representing the *q-gram*.

#### Fundamental Ideas

Let 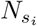 be a set of numbers of a sequence *s*_*i*_, and *π* be a permutation of all the possible integers allowed to appear in such set. The following serves as a *MinHash* signature of *s*_*i*_:

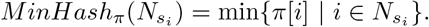

Similar sequences are expected to have the same *MinHash* signature for a given permutation *π* with high probability. Thus, equal signatures may indicate similar sequences. For this reason, it can be concluded that concatenating several *MinHash* signatures for a set of permutations Π, and demanding the two sequences’ signatures to be equal, serve as a reliable approximation of their similarity. The combination of several *MinHash* signatures is referred as an *LSH signature*, as presented in [28]. We denote the length of the signature (the number of *MH* signatures composes it) with *k*.

Another method we used to approximate the distance between two sequences *s*_*i*_, *s*_*j*_ is the Sørensen–Dice coefficient [29], which allows to compare the similarity of 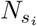 and 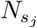. For two vectors *X, Y*, the Sørensen–Dice coefficient is defined by:

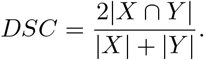

The algorithm requires the *LSH signatures* of sequences to be equal before setting them in the same cluster. We add another condition, as this approach results with many false-positives, and require the Sørensen–Dice similarity to be above a certain threshold.

#### Additional Implementation Details

Unlike the original work in [28], we improve performance by roughly dividing the input into chunks, as shown earlier in Step 1. This process reduces the total number of comparisons needed and minimizes the run time. Moreover, during the run of our algorithm, we also maintain a score for each sequence. The score is a counter, incremented each time a sequence is paired with another. Here we assume that a sequence similar to many other sequences in its cluster (and as a result, has a high score), can serve as a good representative of its cluster in the next step.

#### Parameter Selection

We have control over three parameters - *q* (the size of the *q*-gram), *k* (the length of the *LSH signature*), and the number of iterations of the algorithm. Antkowiak et al. [28] suggest that settings the signature’s length around 3 helps to reflect enough information from the original strand, while still allowing for justified merging decisions later. From similar arguments, a suitable range of values for *Q* would be between 3 and 7. As stated above, the probability of two strands *s*_*i*_ ans *s*_*j*_ to have the same *MinHash* signatures relates to the desired similarity of their number sets. The values used as thresholds for determining similarity are *θ*_*high*_ = 0.32 and *θ*_*low*_ = 0.28. The motivation behind the usage of those values, is an attempt to find a balance between wanting the sequences to be close enough (resulting in similar number sets, and a high Sørensen–Dice similarity), without missing too many related sequences for not being perfectly identical. Following the above, it can be concluded that the probability of two strands with similar number sets having the same signatures is *similarity*^*k*^. When using *k* = 3 such signatures, and the high similarity threshold of *θ*_*high*_ = 0.32, the number of *LSH signatures* that the algorithm needs to compute to ensure finding all pairs of similar strands *s*_*i*_ and *s*_*j*_ with high probability is denoted by: 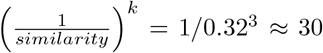. Therefore, we set the required number of iterations to 30.

#### 3.1.3. Step 3 - Full Clustering

The final step is the most dominant step of our clustering process. This step includes iterating over all the clusters created, choosing representatives from each one, and then executing the flow from the previous stage, now in respect to all the sequences without limiting to a certain chunk. This step has proven to be very essential, as it manages to efficiently and accurately handle the numerous single sequences (clusters of size 1) not yet merged into one of the other clusters. As mentioned before, the representatives are selected based on their score. When the number of merges were made during several consecutive iterations has decreased, the representatives are replaced. This follows the fact that other highly scored sequences in a cluster were proved to be helpful as well. The motivation for repeating the same process as in Step 2, but now with the whole input, is the need to merge clusters from different chunks, altogether with better handling the singles. In this step, the algorithm uses two values, namely *θ*_*high*_ = 0.25 and *θ*_*low*_ = 0.22, as thresholds to determine the similarity between clusters. Lower thresholds are used as the algorithm now attempts to handle strands that could not be clustered in the previous step, implying they are likely to be more erroneous.

##### Algorithm 3

Full Clustering

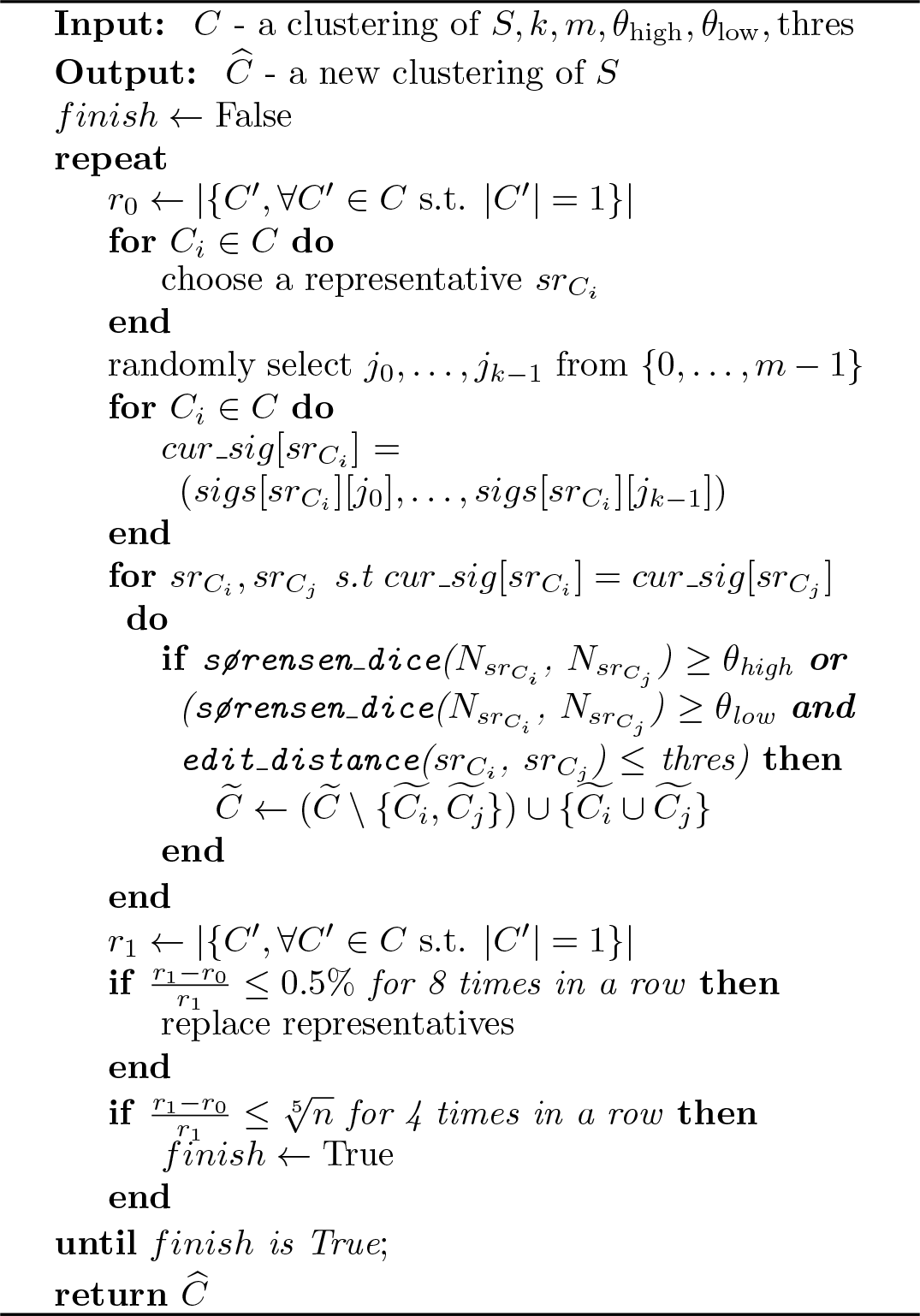

#### 3.1.4. Time Complexity of the Algorithm and Implementation Notes

In this section, we show that the run time for our proposed algorithm is *O*(*nl*) in the worst case, where *n* is the number of sequences in the input, and *l* is the strands length. In the running time analysis of the algorithm, we pay attention to each step on its own.

When splitting the input into chunks in Step 1, the number of iterations is dynamically determined during the run. However, we also use a constant upper bound, not allowing it to exceed a certain number of iterations. The value of this upper bound was deduced empirically. In each iteration, we compare the clusters’ signatures. Both computing the signatures and executing the comparisons is performed in linear time (*O*(*l*)). In practice, using representatives from chunks instead of iterating over all the sequences contributes to a significant performance improvement.

Creating a number set and an *LSH signature* for a single strand depends on its length. Therefore, the setup essential for the entire clustering process in Step 2 is *O*(*nl*). During the clustering process, the sequences are compared using the signatures, but unlike Step 1, an additional check of the similarity is added. Nevertheless, it demands *O*(*l*) operations at most.

The last step is technically identical, aside from refraining from going through the whole input. Instead, we use *O*(1) representatives from each cluster. For this reason, the upper bound remains the same, although the actual run time dramatically improves.

According to the design of the algorithm, several steps can take advantage of multi-core processors. For example, the computation of the number sets and the *LSH signatures* does not require any kind of communication between the cores and can benefit from parallelism. Additionally, in Step 2, the clustering is done in each chunk on its own, suggesting that it is also possible to perform this step in parallel. In our implementation, we support parallel mode which parallelizes some of the steps mentioned above.

## 4. Results

For the testing, a setup with the following specifications was used: 32-cores Intel Xeon E5-2630 CPU, 128GB RAM. Several datasets were tested, with the purpose of evaluating the performance of the algorithm on diverse datasets, with different characteristics. The datasets used differ in the following settings: the number of strands in the original design (equals to the number of clusters in the perfect clustering), the length of the original strands (*l*), the total number of noisy copies in the dataset (*n*), the sizes of the clusters, and the error rates of the technologies used. The datasets that were used consist of simulated datasets, and datasets from prior DNA storage studies [3,4,8,9]. The benchmark analysis was performed with the following clustering algorithms:

1. Rashtchian et al.’s clustering algorithm [27], implemented with 780 serial iterations.
2. The Clover clustering algorithm [25], with its suggested parameters: *HorizontalDrift* = 3, *V erticalDrift* = 3, and *treeDepth* = 15.
3. The LSH-based clustering algorithm presented in [28], with its recommended settings. (*k* = 4, *K*_*LSH*_ = 3).

Starcode [23] is considered to be one of the most appropriate clustering algorithms for DNA storage systems [30]. However, according to Guanjin et al. [25], while Starcode is extremely fast, it fails to achieve high-quality clustering. Furthermore, Rashtchian et al. [27] demonstrated that their clustering algorithm outperforms the Starcode algorithm, despite the fact that the latter was tested using various distance threshold parameters (*d* ∈{2, 4, 6, 8}). Therefore, we have chosen not to include Starcode in our benchmark analysis.

As will be shown later (and will be further discussed in Section 7, the algorithm in [27] exhibits significant instability, as manifested in its correctness and running time. To illustrate this issue, we conducted multiple runs of the algorithm on several datasets, and analyzed the average results and standard deviation values of the metrics. In the remaining datasets, due to time constraints, we present the results obtained from successful runs of the algorithm.

### 4.1. Experimental Datasets

Our experimental datasets were collected from real DNA storage experiments performed in recent years:

I. 2015, Grass et al. [3] - This group encapsulated DNA in an inorganic matrix, and employed error-correcting codes to correct storage-related errors. Specifically, they translated 83KB of information to 4991 DNA strands, each 158 nucleotides long (117 nucleotides of encoded information and 41 nucleotides served as adapters), which were encapsulated in silica.
II. 2017, Erlich and Zielinski [8] - In this work, the researchers encoded 2.11MB of data into 72,000 DNA sequences of length 152. The coding scheme, “DNA Fountain”, was based on the Luby transform with redundancy of 7% (means that 67,088 sequences were information sequences, and the rest were redundancy). The decoding is done with a message-passing decoder (“LDPC”-like). They also applied homopolymer and GC-content constraints in their code.
III. 2018, Organick et al. [4] - They encoded 200MB of data and published one file of 9.5MB of data which was encoded to 607,150 DNA sequences. In their coding scheme they used RS code with 15% redundancy on the symbols. They also used primers to enable random access.
IV. 2021, Yekhanin et al. [9] - The paper introduces a new reconstruction algorithm - Trellis BMA, whose complexity is linear in the number of traces. For their performance comparisons they published a new dataset of 269,709 traces of 10,000 uniform random DNA sequences of length 110 each.

A detailed description of the datasets listed above and their characteristics can be found in Table 1. The evolution of the algorithms on dataset I refers to the results of 10 runs of each algorithm on the dataset. In Figures 1 and 2, it can be clearly seen that our algorithm outperforms all other algorithms. The summary of the average results and the values of the standard deviations between all the runs appear in Tables 2 and 3.

**Table 1:** Properties of experimental datasets.

| Ref. | Data | Copies | Len. | Clust.<br>Sizes | Err. (%) |  |  |
| --- | --- | --- | --- | --- | --- | --- | --- |
|  |  |  |  |  | sub | ins | del |
| [3] | 4,989 | 2,949,757 | 117 | 1-8863 | 0.331 | 0.0503 | 0.604 |
| [8] | 72,000 | 13,33,2276 | 152 | 1-683 | 0.968 | 0.0582 | 0.162 |
| [4] | 596,499 | 11,378,301 | 110 | 1-87 | 0.254 | 0.0412 | 0.134 |
| [9] | 9984 | 269,709 | 110 | 1-164 | 1.07 | 1.64 | 2.01 |

**Table 2:** Results summary of dataset I.

| | Avg. Acc.<br>($\gamma = 0.95$) | Stdev. Acc.<br>($\gamma = 0.95$) | Avg. TS | Stdev. TS |
| --- | --- | --- | --- | --- |
| ours | **0.9991** | 0.0006 | **0.9994** | 0.0007 |
| [27] | 0.7086 | 0.4696 | 0.6926 | 0.4723 |
| [25] | 0.5219 | 0.0058 | 0.9208 | 0.0021 |
| [28] | $2.76 \cdot 10^{-4}$ | $1.84 \cdot 10^{-4}$ | $3.89 \cdot 10^{-4}$ | $3.35 \cdot 10^{-5}$ |

**Table 3:**
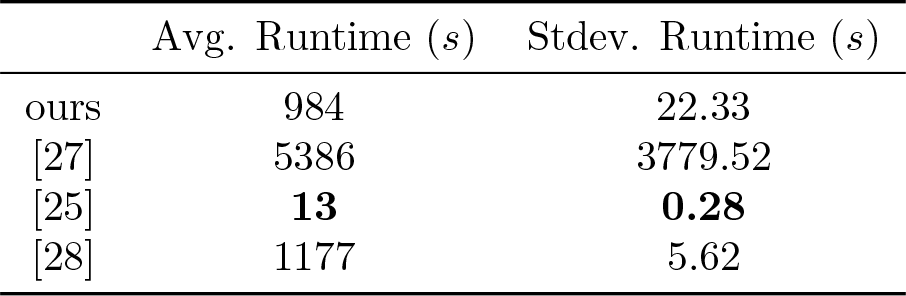
Runtime summary of dataset I.

|  | Avg. Runtime (s) | Stdev. Runtime (s) |
| --- | --- | --- |
| ours | 984 | 22.33 |
| [27] | 5386 | 3779.52 |
| [25] | **13** | **0.28** |
| [28] | 1177 | 5.62 |

**Figure 1.**
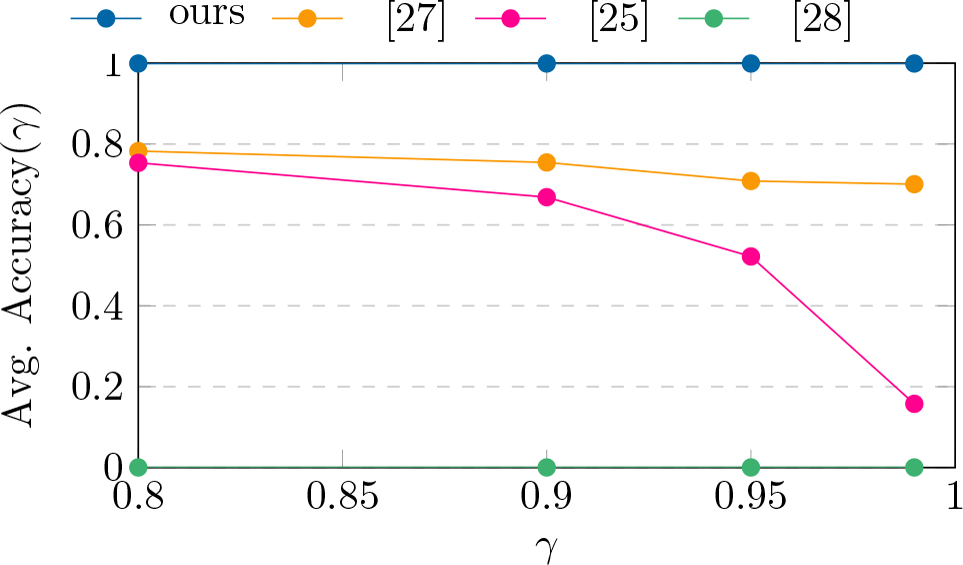
Avg. Accuracy(*γ*) results on dataset I.

**Figure 2.**
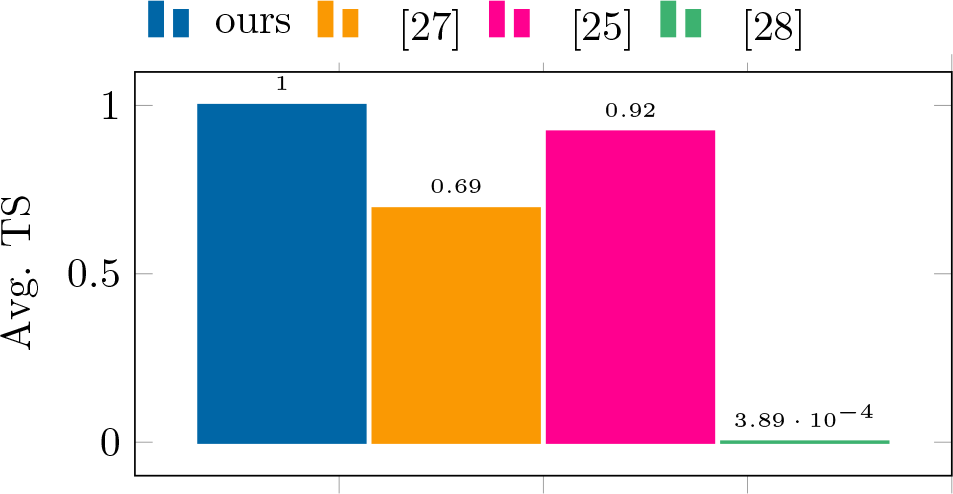
Avg. TS results on dataset I.

Full results of the algorithms on dataset II (for a single successful run) can be found in Figures 10 and 11 at the end of this paper. The TS values of the algorithms and their running times on this dataset can be seen in Figure 9 and Table 7.

Figures 3 and 4 show the performance results of the algorithms (for a single successful run) on datasets III and IV, respectively. Compared to the results obtained from dataset I, the algorithms performed significantly worse. To address this issue, we conducted an additional analysis on these datasets and proposed a solution based on pseudorandomization.

**Figure 3.**
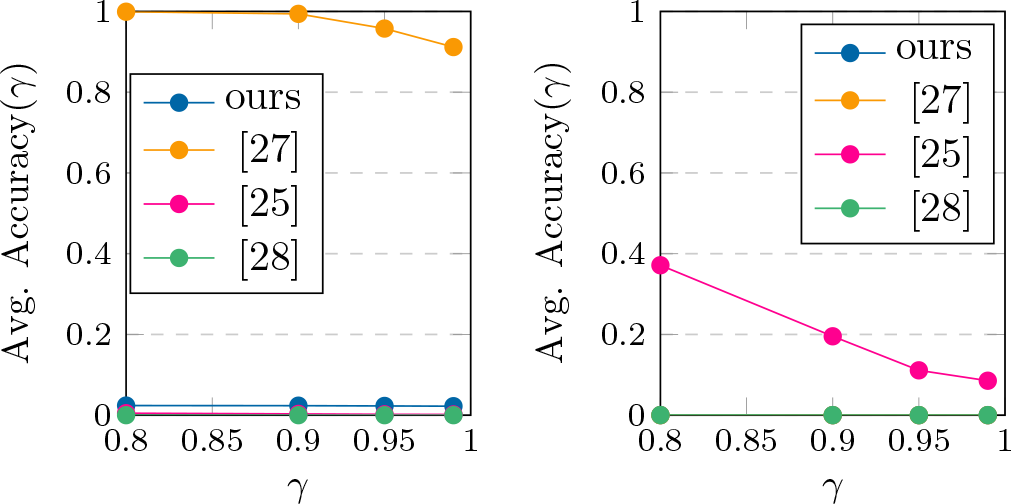
Accuracy(*γ*) results on datasets III and IV.

**Figure 4.**
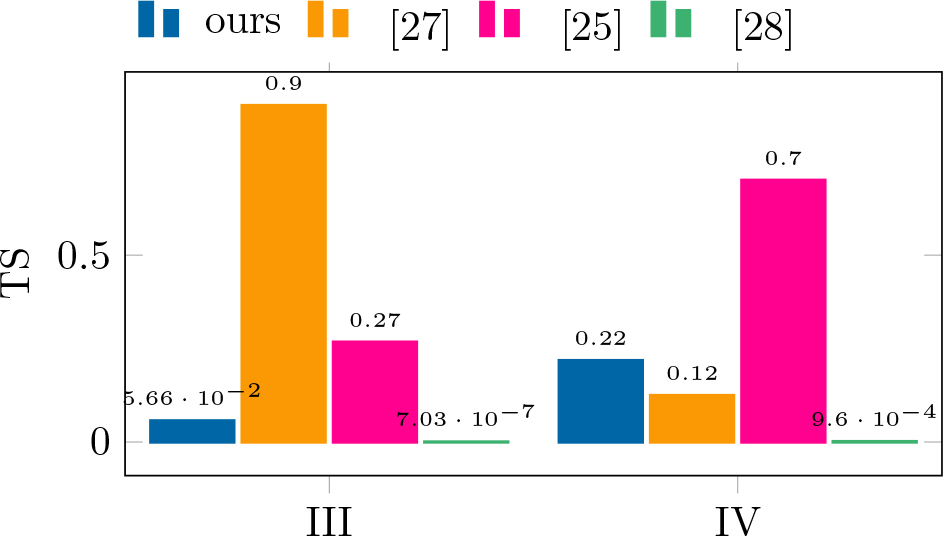
TS results on datasets III and IV.

#### 4.1.1 Enhancing Clustering Performance with Pseudo-Randomized Designs

As mentioned before, all algorithms performed poorly on dataset IV. Moreover, with the exception of [27], all algorithms struggled to cluster dataset III. These results motivated us to conduct further analysis of these two datasets to identify what makes them difficult to cluster. We discovered that in the original design, many long runs of characters appear multiple times in different input strands. These patterns appeared in most of the noisy copies of the input strands. Such patterns have a significant impact on a crucial step in clustering algorithms: the assessment of similarity between two random input strands, particularly when it relies on examining the correlation of their substrings. This critical step enables algorithms to make informed decisions and assign two random input strands to the same original cluster. Therefore, we concluded that such designs are likely to lead clustering algorithms to complete failure.

Here we propose a simple solution for such designs, requiring only a few extra bits of redundancy. Given an input design (with potential similarity among different DNA strands), one can randomly choose a seed and use it to generate pseudo-random DNA strands matching the original design’s length and input set size. Each input strand is then XORed with its corresponding pseudo-random DNA strand, ensuring a high likelihood that the new strands are far from each other (in terms of edit distance) and do not contain repeated substrings across different input strands. To retrieve the original data, pseudo-random strands are regenerated using the original seed and XORed with the received information. The scheme’s redundancy is log(*seed*) = *O*(1), as only extra bits are needed for the seed value.

To verify the correctness and efficiency of this technique, we applied it to the original designs of datasets III and IV. We XOR-wised the original designs with the pseudorandom strands and created clusters with noisy copies for each resulting strand. The cluster sizes were consistent with the original dataset’s cluster sizes. This process was performed using the DNA-Storalator [31] and was based on the error characterization of the original datasets. The results of this scheme are presented in Figures 5 and 6.

**Figure 5.**
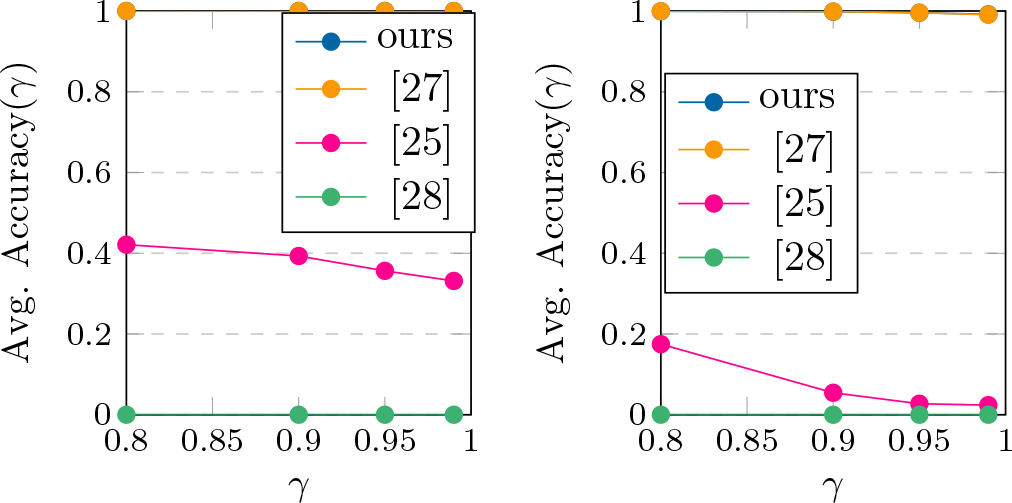
Accuracy(*γ*) results on datasets III^*^ and IV^*^. perturbed datasets.

**Figure 6.**
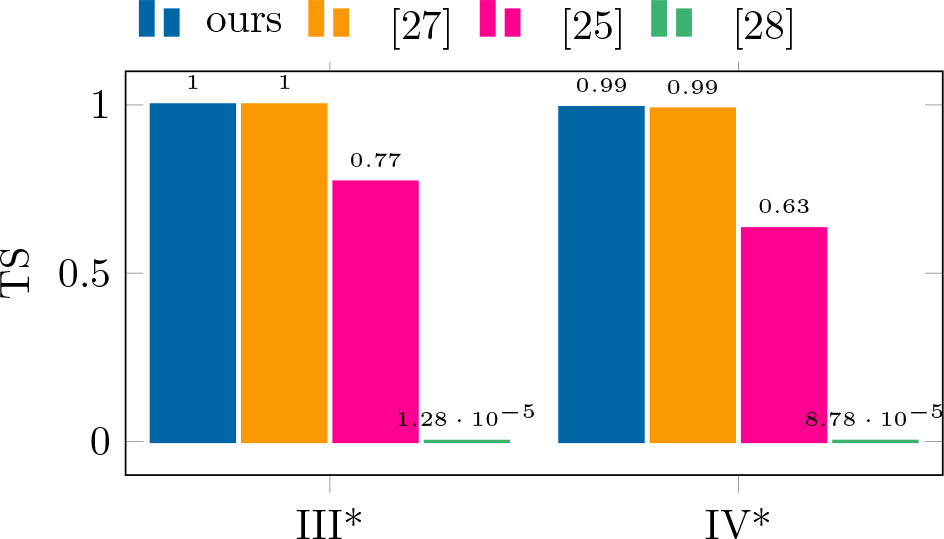
TS results on datasets III^*^ and IV^*^. perturbed datasets.

**Figure 7.**
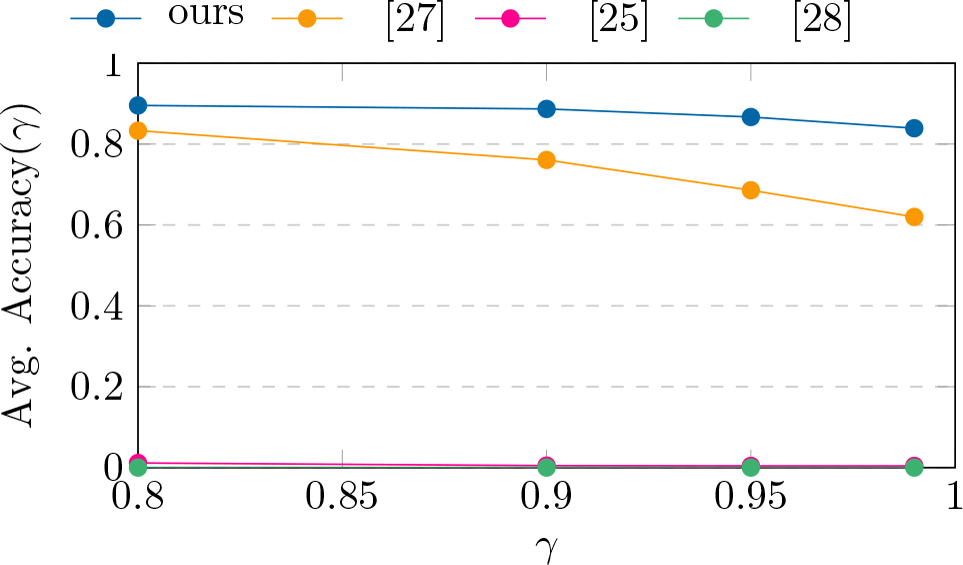
Avg. Accuracy(*γ*) results on dataset VI.

The comparison of the clustering algorithms in Figures 3 and 4 with those in Figures 5 and 6 shows that the pro-posed pseudo-randomization scheme is highly efficient and accurate, leading to a significant improvement in clustering performance. Therefore, it is recommended to use this scheme, either alone or in combination with other encoding methods, to achieve better results in the clustering phase.

### 4.2. Simulated Datasets

All simulated datasets used were generated with uniform random data. The errors were simulated artificially, with values typical for the relevant synthesis technology. The error simulation process was performed using the DNA-Storalator [31], and based on error characterization analysis of the SOLQC tool [32]. Table 3 shows an extended summary of the simulated datasets used and their properties.

**Table 4:** Properties of simulated datasets.

|  | Data | Copies | Len. | Clusters<br>Sizes | Err. (%) |  |  |
| --- | --- | --- | --- | --- | --- | --- | --- |
|  |  |  |  |  | sub | ins | del |
| V | 500,000 | 7,987,578 | 130 | 0-32 | 2.2 | 1.7 | 2 |
| VI | 500,000 | 8,002,591 | 60 | 0-32 | 2.2 | 1.7 | 2 |
| VII | 500,000 | 8,002,736 | 150 | 0-32 | 0.132 | 0.058 | 0.023 |

We randomly selected datasets V and VI to evaluate the performance of the algorithms through multiple runs (10). As shown in Figures 7 and 8, our algorithm outperforms all other algorithms in terms of both TS values and Accuracy(*γ*) index scores on dataset VI. Tables 5 and 6 provide a summary of the performance metrics and running times of the algorithms on this dataset.

**Figure 8.**
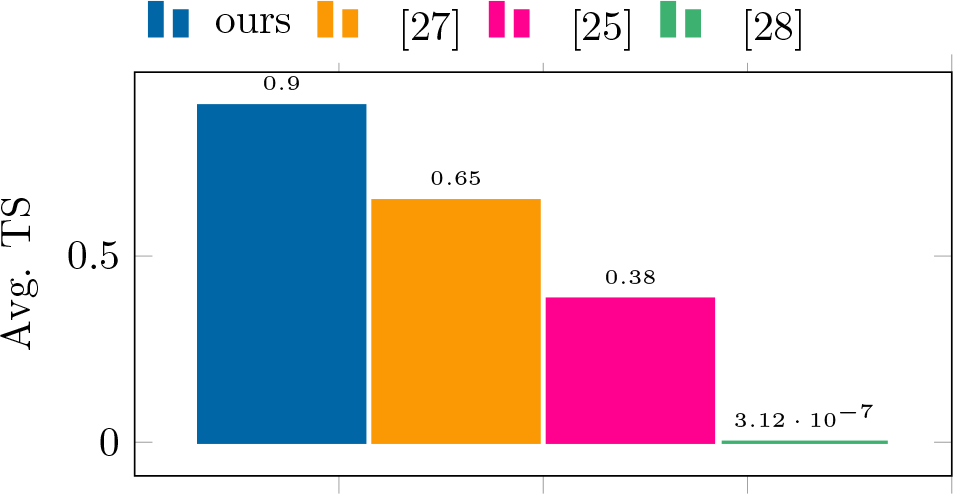
Avg. TS results on dataset VI.

**Figure 9.**
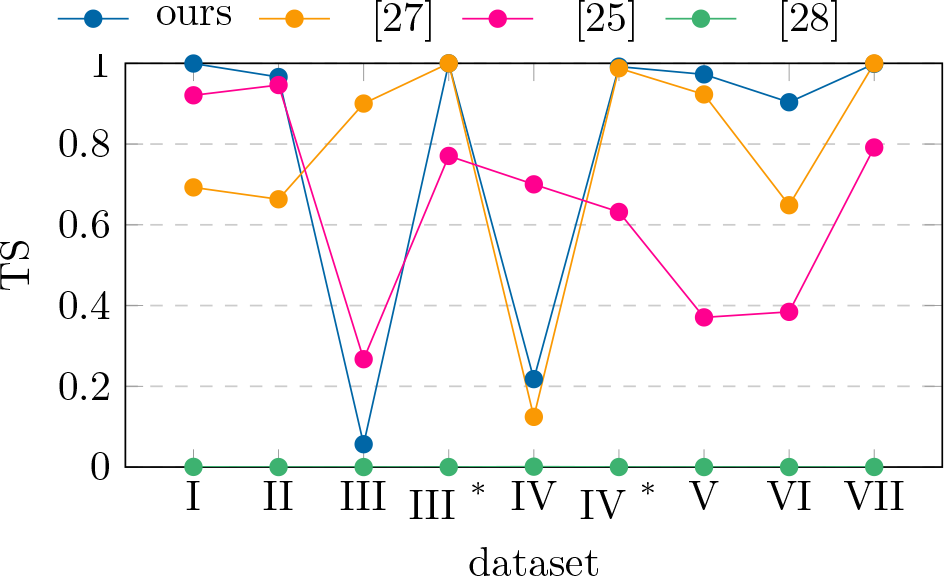
TS results on all datasets.^*^ perturbed datasets

**Figure 10.**
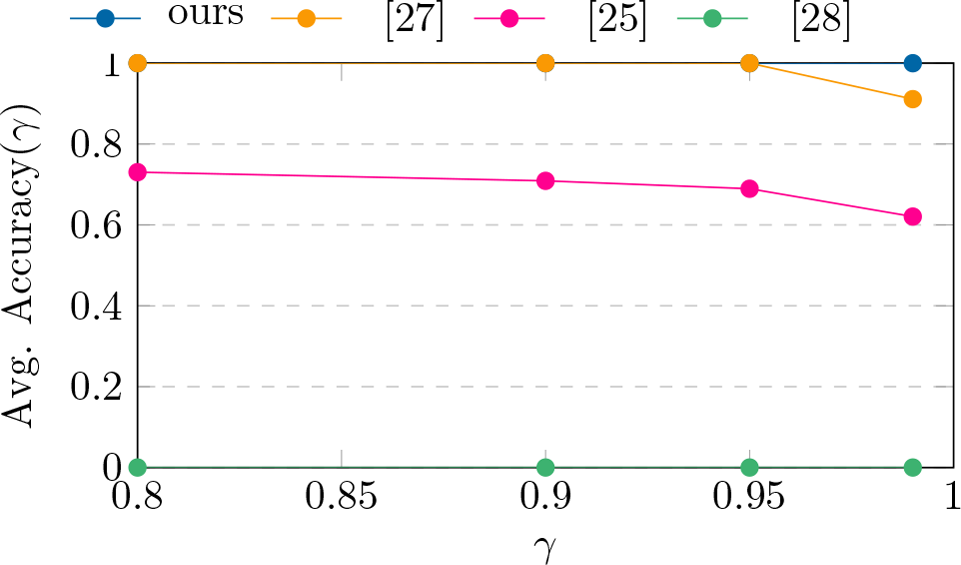
Accuracy(*γ*) results on dataset II.

**Figure 11.**
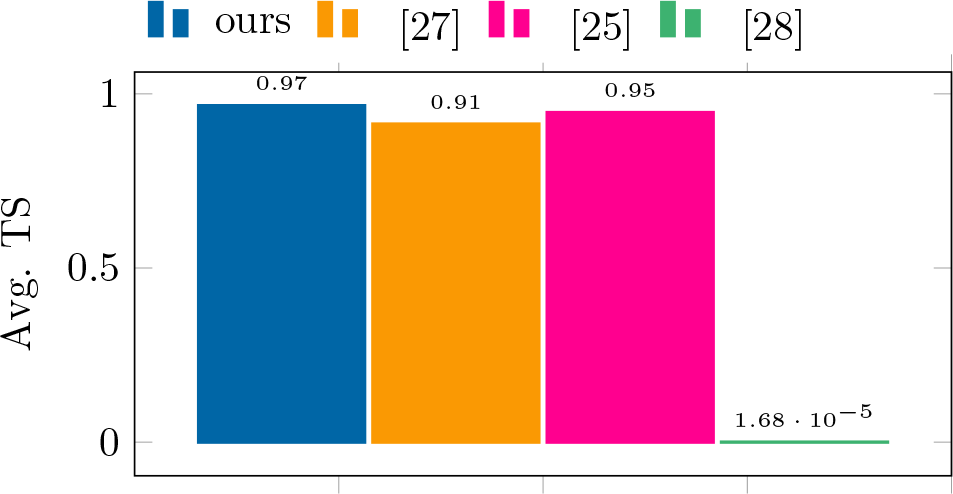
TS results on dataset II.

Full results of the algorithms on datasets V and VII can be found in Figures 12-15 and Tables 8-9 at the end of this paper, as well as in the Results Summary.

**Table 5:** Results summary of dataset VI.

| | Avg. Acc.<br>($\gamma = 0.95$) | Stdev. Acc.<br>($\gamma = 0.95$) | Avg. TS | Stdev. TS |
| --- | --- | --- | --- | --- |
| ours | **0.8669** | 0.0059 | **0.9036** | 0.0042 |
| [27] | 0.6858 | 0.3176 | 0.6485 | 0.2808 |
| [25] | 0.0042 | $8.66 \cdot 10^{-5}$ | 0.3843 | 0.0002 |
| [28] | 0 | 0 | $3.12 \cdot 10^{-7}$ | 0 |

**Table 6:** Runtime summary of dataset VI.

|  | Avg. Runtime (s) | Stdev. Runtime (s) |
| --- | --- | --- |
| ours | 9934 | 770.85 |
| [27] | 13525 | 9270.39 |
| [25] | **758** | **8.98** |
| [28] | 3410 | 71.08 |

**Table 7:** Runtime summary of all datasets (*s*). perturbed datasets, ^**^ average running times, ^***^ serial runs (due to memory limitations).

| dataset | ours | [27] | [25] | [28] |
| --- | --- | --- | --- | --- |
| I | 2440** | 7538** | **47\*\*** | 1177** |
| II | 21781*** | 13280 | **265** | 6119 |
| III | 17125*** | 11856 | **1246** | 4796 |
| III* | 16598*** | 9989 | **671** | 5111 |
| IV | 168 | 208 | **18** | 102 |
| IV* | 155 | 317 | **18** | 104 |
| V | 12946** | 18783** | **776\*\*** | 3513** |
| VI | 9934** | 13525** | **758\*\*** | 3410** |
| VII | 11637 | 14307 | **554** | 3444 |

**Table 8:** Results summary of dataset V.

| | Avg. Acc.<br>($\gamma = 0.95$) | Stdev. Acc.<br>($\gamma = 0.95$) | Avg. TS | Stdev. TS |
| --- | --- | --- | --- | --- |
| ours | **0.9809** | 0.0017 | **0.9726** | 0.0022 |
| [27] | 0.9475 | 0.0919 | 0.9229 | 0.1279 |
| [25] | 0.0039 | $9.45 \cdot 10^{-05}$ | 0.3707 | 0.0003 |
| [28] | 0 | 0 | $1.95 \cdot 10^{-6}$ | 0 |

**Table 9:**
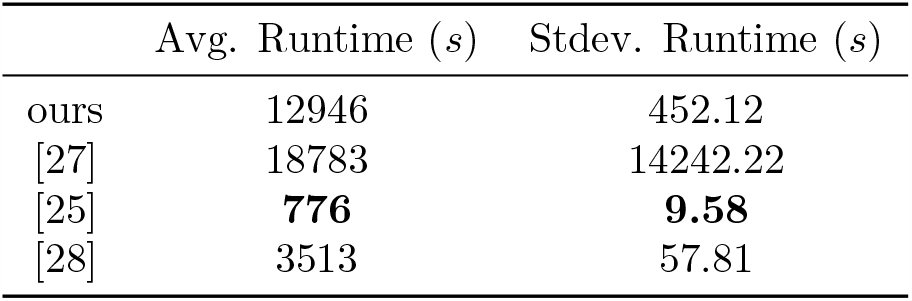
Runtime summary of dataset V.

|  | Avg. Runtime (s) | Stdev. Runtime (s) |
| --- | --- | --- |
| ours | 12946 | 452.12 |
| [27] | 18783 | 14242.22 |
| [25] | **776** | **9.58** |
| [28] | 3513 | 57.81 |

**Figure 12.**
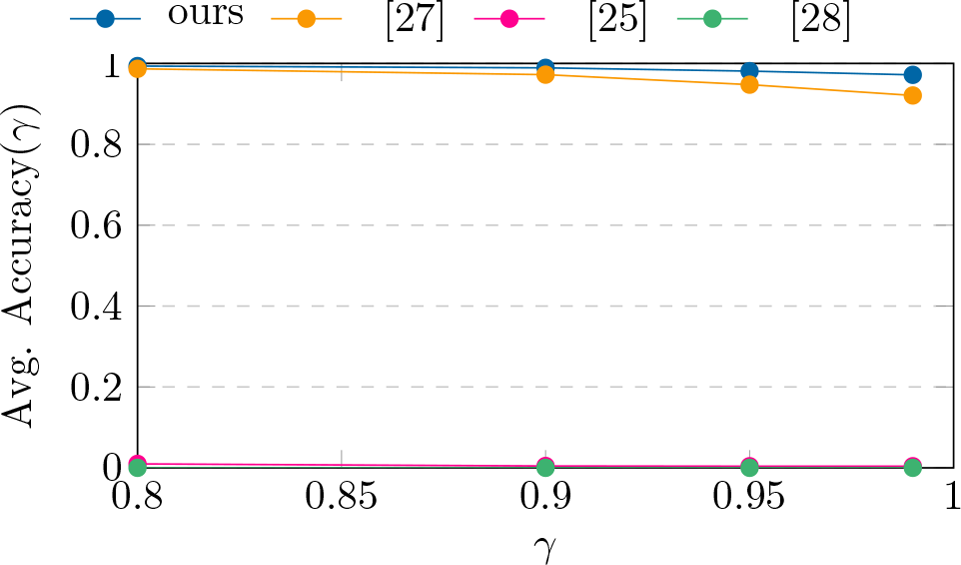
Avg. Accuracy(*γ*) results on dataset V.

**Figure 13.**
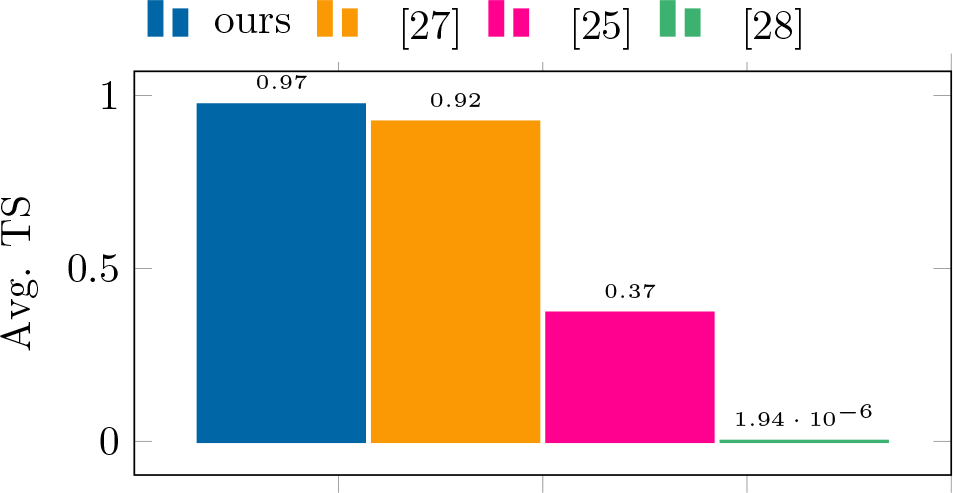
Avg. TS results on dataset V.

**Figure 14.**
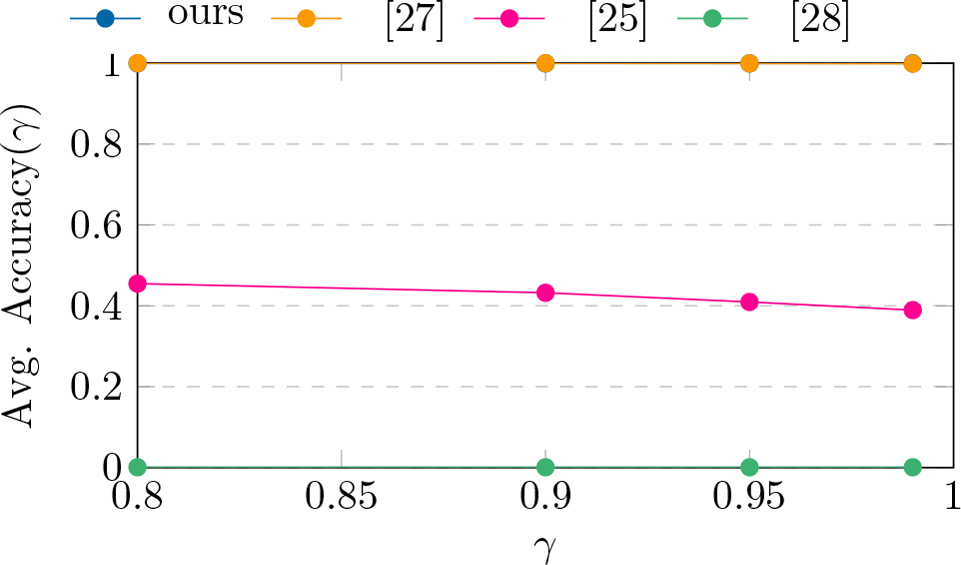
Accuracy(*γ*) results on dataset VII.

**Figure 15.**
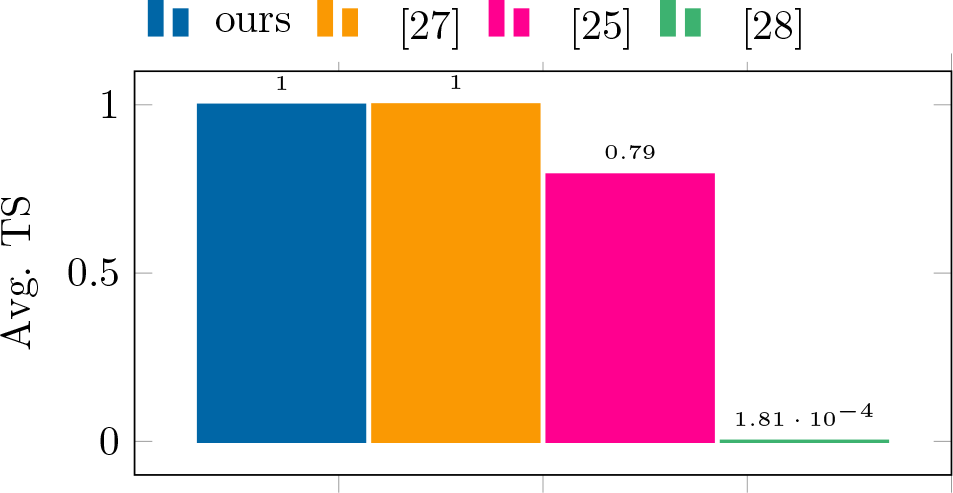
TS results on dataset VII.

#### Overall Results Summary

Figure 9 summarizes a comprehensive overview of the TS results for all datasets, highlighting trends and enabling clear comparisons between the general correctness of all the algorithms. Table 7 summarizes the running times of the algorithms. dataset

## 5. Availability and Implementation

GradHC is freely available at https://github.com/bensdvir/GradHC. Our implementation uses Python and includes both serial and parallel modes.

## 6. Supplementary information

Access to all datasets (both simulated and experimental) will be provided upon request.

## 7. Discussion

The algorithms that produced the best results on all tested datasets are ours and the clustering algorithm in [27]. As expected, better clustering performance was achieved by all algorithms as the error rates decreased. In terms of runtime, Clover [25] emerged as the fastest clustering algorithm, outperforming all other algorithms by several orders of magnitude. Clover achieves its speed by constructing a tree structure to search for a specified interval, instead of computing the Levenshtein distance between strands. However, despite its impressive speed, Clover was only able to achieve high-quality clustering results on datasets I and II, which are characterized by relatively low error rates and very large clusters. Furthermore, due to complexity and slowness of the synthesis and sequencing processes involved in DNA-based storage, it is mainly intended for long-term storage such as archives [2, 3]. Thus, clustering speed is not the highest priority in such systems. In contrast, the LSH-based clustering algorithm [28] showed poor performance on all tested datasets, both simulated and experimental. It is important to note that although the algorithm’s results were unsatisfactory, Antkowiak et al. [28] were able to recover all the information encoded in the experiment by applying a coarse filtering on the clusters created and by using an outer code that could handle a high percentage of missing strands.

During the benchmark performance, we observed that the implementation of [27] often exhibited unexpected behavior, whereas our algorithm and all other suggestions remained stable. To demonstrate the weakness of the algorithm in [27], we conducted multiple runs of the algorithm on datasets I, V, and VI (selected arbitrarily). This observation is reflected in the algorithm’s TS and Accuracy(*γ*) standard deviation values, as shown in Tables 2 and 5. As can be seen, the standard deviation values of the clustering algorithm in [27] are several orders of magnitude greater than the standard deviation values of the other algorithms, which close to 0. Additionally, when the algorithm fails to converge, its instability can significantly affect its running time. This is evident in the high standard deviation values of the algorithm’s runtime (see Tables 3 and 6).

Occasionally, the clustering algorithm suggested in [27] failed to converge, impacting both the runtime and the metrics used to evaluate the measurements. Our hypothesis is that datasets containing small clusters may cause a situation where a series of incorrect merges in the early stages is sufficient for hurting the rest of the process. The reason originates from the fact that not all the strands participate in every iteration, and only representatives (whose selection is random) are used, making the algorithm vulnerable if poor decisions were made at the start. A key feature of our proposal appears in Step 2, in which the clustering process is done with the majority of the sequences taking a part, instead of picking representatives. In addition, when representatives are used in Step 3, they are selected intelligently, according to their score. In this manner, we were able to overcome scenarios that could have failed the algorithm presented in [27].

In conclusion, the most significant advantage of our proposed approach is its ability to deliver consistent high-quality results, even when processing challenging inputs, such as short designs, small clusters, and high error rates. Although the algorithm in [27] can produce results (in terms of both runtime and correctness) that are comparable to or better than those of our algorithm in multiple runs, it often fails to provide satisfactory outcomes, as evidenced by the standard deviation values obtained from multiple runs of the algorithm. In such scenarios, it is highly probable that a DNA storage system relying on an algorithm whose convergence cannot be guaranteed would fail to retrieve the stored information.

## Footnotes

1 A string metric allows for quantifying how similar two strings are to each other. It is measured by counting the minimum number of operations required to transform one string into the other.

2 Minimum number of single-character edits (insertions, deletions or substitutions) required to change one word into the other.

